# Galectin-1 promotes the invasion of bladder cancer urothelia through their matrix milieu

**DOI:** 10.1101/441642

**Authors:** A. Balakrishnan, D Pally, K. Gondkar, S. Naito, D. Sidransky, A. Chatterjee, P. Kumar, R. Bhat

## Abstract

The progression of carcinoma of the urinary bladder involves migration of cancer epithelia through their surrounding tissue matrix microenvironment. This was experimentally confirmed when a gender- and grade-diverse set of bladder cancer cell lines were cultured in pathomimetic three-dimensional laminin-rich environments. The high-grade cells, particularly female, formed multicellular invasive morphologies in 3D. In comparison, low- and intermediate-grade counterparts showed growth-restricted phenotypes. A proteomic approach combining mass spectrometry and bioinformatics analysis identified the estrogen-driven lactose-binding lectin Galectin-1 (GAL-1) as a putative candidate that could drive this invasion. Expression of *LGALS1*, the gene encoding GAL-1 showed an association with tumor grade progression in bladder cell lines. Immunohisto- and cyto-chemical experiments suggested greater extracellular levels of GAL-1 in 3D cultures of high-grade bladder cells and cancer tissues. High levels of GAL-1 associated with increased proliferation- and adhesion- of bladder cancer cells when grown on laminin-rich matrices. Pharmacological inhibition and Gal-1 knockdown in high-grade female cells decreased their adhesion to, and viability on, laminin-rich substrata. Higher GAL-1 also correlated with reduced E-cadherin and increased N-cadherin levels in consonance with a mesenchymal-like phenotype that we observed in 3D culture. The inhibition of GAL-1 reversed the stellate invasive phenotype to a more growth-restricted one in high-grade cells embedded within both basement-membrane-like and stromal collagenous matrix scaffolds. Finally, inhibition of GAL-1 specifically altered cell surface sialic acids, suggesting the mechanism by which the levels of GAL-1 may underlie the aggression and poor prognosis of invasive bladder cancer, especially in women.

## Introduction

Urothelial carcinoma of the bladder (UCB) is a heterogeneous disease and its management is challenging because bladder tumors frequently recur (1). The rate of recurrence of these tumors has been reported to be due to the result of ‘field effect’ and/or residual presence of tumor cells following surgery (2). Although men are about 3 to 4 times more likely to develop bladder cancer during their lifetime than women, the latter often present with more advanced cancers resulting in poorer survival and therapeutic outcomes (3).

Based on the extent of invasion Bladder tumors may either be classified as non-muscle-invasive (NMIBC) (Ta/T1), and muscle-invasive (MI) (T2/T3). At diagnosis, the majority of bladder cancers are non-muscle-invasive (stage Ta) tumors of low grade. NMIBC frequently recur (50-70%) and only 10-15% progress to invasion. On the other hand, MI tumors are diagnosed de novo and have poorer prognosis (1). The intra- and inter- tumor heterogeneity of this cancer does not allow it to be prognosed by a single biomarker. Tremendous efforts have been put in identifying a robust set of biomarkers for early diagnosis, patient stratification, treatment selection and outcome and prognosis. However, the highly heterogeneous nature of these tumors precludes success in the field. A previous study from our group had identified multiplex protein signatures for the detection of bladder cancer in voided urine samples. Our biomarker panel consists of 5 robust biomarkers offers unprecedented accuracy for the diagnosis of bladder cancer patients (4). However, the oligoclonal nature of these tumors where a single clone spreads via intraepithelial migration could also contribute to the heterogeneity of individual bladder cancer. In comparison with advances on the diagnosis of bladder cancer, our understanding of bladder cancer cell migration through the organ stroma during metastasis is still inadequate. Key signaling pathways such as NF-κB and TGFβ have been implicated in the induction of bladder cancer cell migration. Upregulation of the p65, which is involved in heterodimerization of NF-κB, results in movement of cancer cells through the ubiquitinylation and degradation of RhoGDIα (5). Ablation of TGFβ within a murine model of bladder carcinoma results in poorer invasion and progression of bladder cancer cells along with a concomitant downregulation of mesenchymal markers (and upregulation of epithelial counterparts) (6). Interestingly, the levels of a non-canonical IkB kinase, TBK1 have been shown to be upregulated in bladder cancer: pharmacological inhibition of its kinase activity attenuates transwell migration of bladder cancer cells, through a decreased phosphorylation of Akt (7). In addition, the upregulation of miR-1-3p has been shown to attenuate the proliferation and migration of cancer epithelia through upregulation of the soluble protein SFRP-1 (8). MiR135a on the other hand, increases proliferation and invasion in a Wnt signaling-dependent manner (9). These studies have predominantly focused on the signaling dynamics associated with cancer cell migration, which is experimentally investigated through movement of cells across permeable filtration barriers. Invasion seen in solid tumors in vivo involves cells navigating through, and interacting with, their stromal matrix milieu (10); there is scarce information on how the proteins and sugars on the surface of bladder cancer cells may interact with bladder ECM in order to migrate to the outer muscle layers.

Cell line models offer the opportunity to assess and recapitulate the biology of neoplastic cells. We have recently performed MS-based quantitative proteomic profiling (data not published) of 7 well-characterized neoplastic bladder cell lines (KK47, SW780, RT112, VMCUB-1, T24, J82, and UMUC3), which were matched with tumor grade (G1=KK47, SW780, G2=RT112, VM-CUB-1 and G3=T24, J82) and also grouped into male (KK47, VM-CUB-1, J82) and female (SW780, RT112, T24) cell types. A prominent dysregulated molecule that features within our analysis is Galectin-1 (GAL-1). This observation is congruent with increasing evidence of aberrant cancer cell-surface glycosylations, and cognate changes in lectins: proteins that sense, interact, and signal through, these aberrant glycans (11). The association of one such galactose-binding lectin: GAL-1 with cancer progression has been widely reported in different tumors type (12–17). Studies have shown that GAL-1 de-regulation affects cell proliferation, adhesion and invasiveness in various tumor types through a host of mechanisms that is dependent on its localization. In the cytoplasm it can interact with oncogenic H-Ras to mediate cell transformation (18). GAL-1 expression in non-malignant developing mammary epithelia and in breast cancer cells in 3D known to localize within nucleus and induce the expression of genes that code for the expression of proliferation- and invasion-specific proteins (12). Notably, GAL-1 has also been identified as for a regulator of pre-mRNA splicing (19,20). Shen and coworkers showed that the elevated levels of GAL-1 increases proliferation and migration of bladder cancer cells through upregulation of MMP-9 in a JNK-dependent manner (21). The gene encoding GAL-1, i. e., *LGALS1* is known to be regulated by estrogen (22). In this manuscript, we ask whether elevated levels of GAL-1 expression correlates with a greater invasion of cancer cells through the stromal microenvironment of the bladder. The altered expression of GAL-1 and the gender-specific disparities could be used as a prognostic marker and a possible target for bladder cancer therapy.

## MATERIALS AND METHODS

### Reagents

All of the chemical reagents were of analytical grade, obtained from commercial suppliers, and used without further purification. The antibodies and reagents used in the study along with their source is mentioned as follows: rabbit polyclonal α-Gal-1 antibody (Abcam ab25138) (used for western blotting and IHC), mouse monoclonal α-Gal-1 antibody (sc-271819, Santa Cruz Biotechnology, USA) (used for ICC), rabbit polyclonal N-Cadherin antibody (#4061, Cell Signalling Technology, USA), rabbit monoclonal E-Cadherin antibody (#3195, Cell Signalling Technology, USA), rabbit polyclonal GAPDH antibody (CSB-PA00025A0Rb, Cusabio, USA), Dulbecco’s modified Eagle medium (DMEM) (HiMedia, India), Fetal Bovine Serum (FBS) (Life Technologies, USA), penicillin-streptomycin (HiMedia, India), Matrigel™ (Corning Life Sciences, USA), trypsin (HiMedia, India), paraformaldehyde (Merck, India), Triton X-100 (HiMedia, India), anti-mouse antibody conjugated with Alexa fluor488 (Invitrogen, USA), DAPI (4′,6-Diamidine-2′-phenylindole dihydrochloride) (Invitrogen, USA), Phalloidin conjugated with Alexa fluor 568 (Invitrogen, USA), BSA (HiMedia, India), collagen (Gibco, USA), Rezazurin sodium salt powder (Sigma Aldrich, India), Propidium iodide (HiMedia, India), TRIzol™ reagent (Invitrogen, USA), DAB substrate (Thermo Fischer Scientific, USA), Hematoxylin (HiMedia, India), Turbofect (Thermo Fischer Scientific, USA), MTT [3-(4,5-dimethylthiazol-2-yl)-2,5-diphenyltetrazolium bromide] (HiMedia, India) and DMSO (Dimethyl sulfoxide) (Merck, India)

### Cell culture

The following cell lines were used in the study: SW780, RT112, T24, KK47, VM-CUB-1 and J82. All the cell lines except KK47 were purchased from ATCC. KK47 was obtained from Dr. Seiji Naito, Kyushu University, Fukuoka 819-0935, Japan. The cells are maintained in Dulbecco’s modified Eagle medium (DMEM) supplemented with 10% fetal bovine serum (FBS) and 1X penicillin-streptomycin. The three-dimensional laminin-rich extracellular matrix (lrECM) on-top and embedded cultures were prepared by seeding the trypsinized cells over and within a thin layer of Engelbreth-Holm-Swarm tumor extract-Matrigel™. Three-dimensional Type-1 collagen scaffolds were prepared by adding 8 volumes of acid extracted unpolymerized Type 1 collagen (Gibco) along with 1 volume of 10X DMEM and an appropriate volume of 0.1 N sodium hydroxide to bring the final concentration of polymerizing collagen to 1 mg/ml (pH: 7.4). The scaffolds were polymerized by incubating at 37°C for 30 minutes. Subsequently trypsinized cells were added on top of the scaffolds and the cultured for 3-4 days.

### Immunocytochemistry

The cells from 3D lrECM cultures were pre-treated with 18% and 30% sucrose before fixing with 4 % paraformaldehyde in PBS. For 2D-and collagen cultures, the cells were directly fixed with 4 % paraformaldehyde. The cells were then permeabilized with 0.1 % Triton X-100 in PBS; blocked with PBS containing 3% BSA and 0.1% Triton X-100 for 1 hour in room temperature; followed by primary antibody incubation (α-Gal-1 antibody overnight at 4°C. The cells are then subjected to secondary antibody staining (anti-mouse antibody conjugated with Alexa Fluor 488, DAPI (4′,6-Diamidine-2′-phenylindole dihydrochloride) and Phalloidin conjugated with Alexa Fluor 568. The images of cells plated as monolayers were acquired using an Olympus IX71 microscope and analysed by ImageJ. The images of 3D cultures were obtained by laser scanning confocal microscopy using a Zeiss LSM880 (Airyscan) and analysed by ZEN Lite software.

### Lectin cytochemistry

The cells were fixed with 4% paraformaldehyde and blocked with PBS containing 3% BSA for 1 hour. Cultures were incubated in 1:1000 of FITC-conjugated *Limulus polyphemus* Lectin (LPA) (EY laboratories, Inc., USA), *Aleuria Aurantia* lectin (AAL) (Vector laboratories, USA), Jacalin, *Phaseolus vulgaris Erythroagglutinin* (PHA-E) and *Phaseolus vulgaris Leucoagglutinin* (PHA-L) for 3 hours in room temperature. Counterstaining was done using DAPI and Phalloidin conjugated with Alexa Fluor-568. The fluorescent images were obtained in Olympus IX71 microscope and analysed by ImageJ. The corrected total cell fluorescence (CTCF) value was calculated for each cell by the following formula.

CTCF = Integrated Density − (Area of selected cell X Mean fluorescence of background readings)

### Western blotting

The cells were grown in tissue culture dish to 75% confluence and washed with ice cold PBS thrice. Then the cells were treated with RIPA lysis buffer (0.1% SDS, 50mM Tris HCl (pH 7.4), 150mM NaCl, 1% sodium deoxycholate, 1% Nonidet P-40, 1% Triton X-100 with protease inhibitor cocktails). The protein concentration was measured by DC protein assay (Bio-Rad). 30 μg of the protein lysate was resolved by SDS-PAGE and transferred onto methanol pre-treated PVDF (Polyvinylidene difluoride) membrane. The membrane was blocked with 5% fat free milk in TBST (Tris-buffered saline + 0.1% Tween 20) for 15 minutes and then incubated with the primary anti-body (α-Gal-1 antibody overnight at 4°C. It is then treated with secondary antibody (anti-rabbit IgG conjugated with HRP). The proteins were then visualized using chemiluminescence (ThermoFischer Scientific, USA).

### RT-qPCR

RNA from the 6 cell lines (SW780, RT112, T24, KK47, VM-CUB-1 and J82) was isolated using TRIzol™ reagent as per manufacturer protocol. Isolated RNA was quantified using UV-visible spectrophotometer (NanoDrop™). RNA (1μg) was reverse transcribed using Verso™ cDNA synthesis kit (Thermo Scientific AB-1453). All sample were processed at the same time and resulting cDNA was diluted 1:10. Real time PCR with SYBR green detection system (Thermo Fischer Scientific) was performed using StepOnePlus™ real-time PCR system (ABI) and *LGALS1* primers: Forward 5′-TCAAACCTGGAGAGTGCCTT-3′ and Reverse 5′-CACACCTCTGCAACACTTCC-3′ (Annealing temperature: 60°C). Appropriate no-RT and no template controls were included in each biological repeat.

### Immunohistochemistry

The tissues sections were made from paraffin embedded blocks of bladder cancer patients from Kidwai Memorial Institute of Oncology, Bengaluru with the Institutional Human Ethics Committee approval and informed consent of the patients. The slides were first deparaffinised and then subjected to antigen retrieval in citrate buffer. Thereafter, the tissue sections were quenched using hydrogen peroxidase for 30 minutes and incubated with primary antibody (α-Gal-1 antibody overnight at 4°C. The sections were then treated with secondary antibody (anti-rabbit IgG conjugated with HRP). The color was then developed using DAB substrate and counterstained with hematoxylin.

### Adhesion assay

96 well plate was coated with a thin layer (100 μg/ml) of Matrigel ™ in PBS and was allowed to solidify overnight. A total of 5×10^3^ cells suspended in DMEM/ supplemented with 10% FBS was added to each well and incubated for 30 minutes at 37°C in CO_2_ incubator. The cells were then fixed with 4% paraformaldehyde and stained with propidium iodide (50 μg/ml) and the fluorescence (535/617 nm) was measured using a plate reader (Tecan infinite M200 Pro™).

### Resazurin assay

5×10^3^ cells were seeded in each of the 96 well plate and was serum-starved overnight. The cells were then cultured in DMEM (10% FBS) for 24 hours and then treated with resazurin (100 μg/ml) at 37°C in CO_2_ incubator for approximately 1 hour. The cellular proliferation is then assayed by measuring fluorescence at 560/590 nm (Tecan infinite M200 Pro™).

### Genetic perturbation of *LGALS1*

The shRNA clone used to knock out *LGALS1* was 5′-CCGGCCT GAAT CT CAAACCT GGAG ACT CGAGT CTCCAGGTTT GAGATT CAGGTTTTT G 3′ obtained from MISSION shRNA library (TRCN0000057425, Sigma Merck, USA). The plasmid containing the shRNA was packed into lentivirus by packaging vectors (pMD2.G and psPAX2, Addgene, USA (kind gift from Prof. Deepak K Saini)). The plasmids were transfected into 293FT cells (Invitrogen, USA) by Turbofect and cultured in DMEM containing 10% FBS. After 48 hours, the culture medium was filtered using 0.2 μm filter, concentrated using Lenti-X Concentrator (Takara Bio Inc., Japan) and stored at −80 °C. To transduce T24 cells, 2 × 10^4^ cells were cultured in 12 well plate along with lentivirus and polybrene (0.8 μg/ml) for 24 hours. The *LGALS1* knock down cells were selected using puromycin (2 μg/ml).

### OTX008 treatment

OTX008 was supplied from Cayman chemicals (CAS no 286936-40-1) and was dissolved in DMSO (stock 10 mM), further diluted in culture medium before *in vitro* assay.

### MTT assay

The IC_50_ of T24 cells with OTX008 was determined with MTT [3-(4,5-dimethylthiazol-2-yl)-2,5-diphenyltetrazolium bromide] assay. 5 × 10^3^ cells were seeded in 96 well plate and the cells were treated with appropriate drug concentrations for 24 hours. The formazan crystals, which were formed after the addition of MTT solution were dissolved using DMSO and absorbance was measured at 570 nm using a plate reader (Tecan infinite M200 Pro™).

### Statistics

The numerical values of all the experiments were recorded in Excel (Microsoft) and statistical analyses were performed using Graphpad Prism (GraphPad Software, Inc.). The data were analysed using unpaired student’s t test or one-way ANOVA.

## RESULTS

### Grade 3 female bladder cell lines exhibit maximal invasion within 3D basement membrane-like microenvironment

In order to verify if female bladder cancer urothelial grow more aggressively, a panel of six bladder cancer lines distinct in the gender and grade of source tissue were cultured,: the cell lines tested were SW780 (Female; Grade 1) (23), RT112 (Female; Grade 2) (24), T24 (Female; Grade 3) (25), KK47 (Male; Grade 1) (26–29), VM-CUB-1 (Male; Grade 2) (23,26) and J82 (Male Grade 3) (23,26,30,31). All six cell lines were grown either embedded within, or on top of, laminin-rich extracellular matrix (lrECM), fixed and stained for F-actin (using Phalloidin conjugated with Alexa 568) and DNA (using DAPI). The morphologies were compared with each other and characterized on the basis of phenotypic descriptors used to describe a diverse set of breast cell line cultures by Kenny and coworkers (32) (Fig. 1). Irrespective of gender, low-grade cells (SW780, KK47) organized into spherical or “round” growth-restricted structures in lrECM. Both male and female intermediate-grade cancer cells (RT112, VM-CUB-1) formed “grape” like restricted morphologies. Male high-grade cells (J82) also organized into grape-like structures with mononuclear extensions at the edge of each multicellular cluster. Despite the invasion, the clusters remained discrete within the matrix environment. In contrast, female high-grade cells (T24) proliferated and invaded through the whole scaffold to such extents that independent clusters were bridged by long collective multicellular migrations. This morphology resembles the invasion of mesenchymal invasive breast cancer cells such as MDA-MB-231 in 3D. Our observations suggested that both gender and grade might contribute to the aggressive phenotype of T24 cells in 3D.

**Figure 1:**
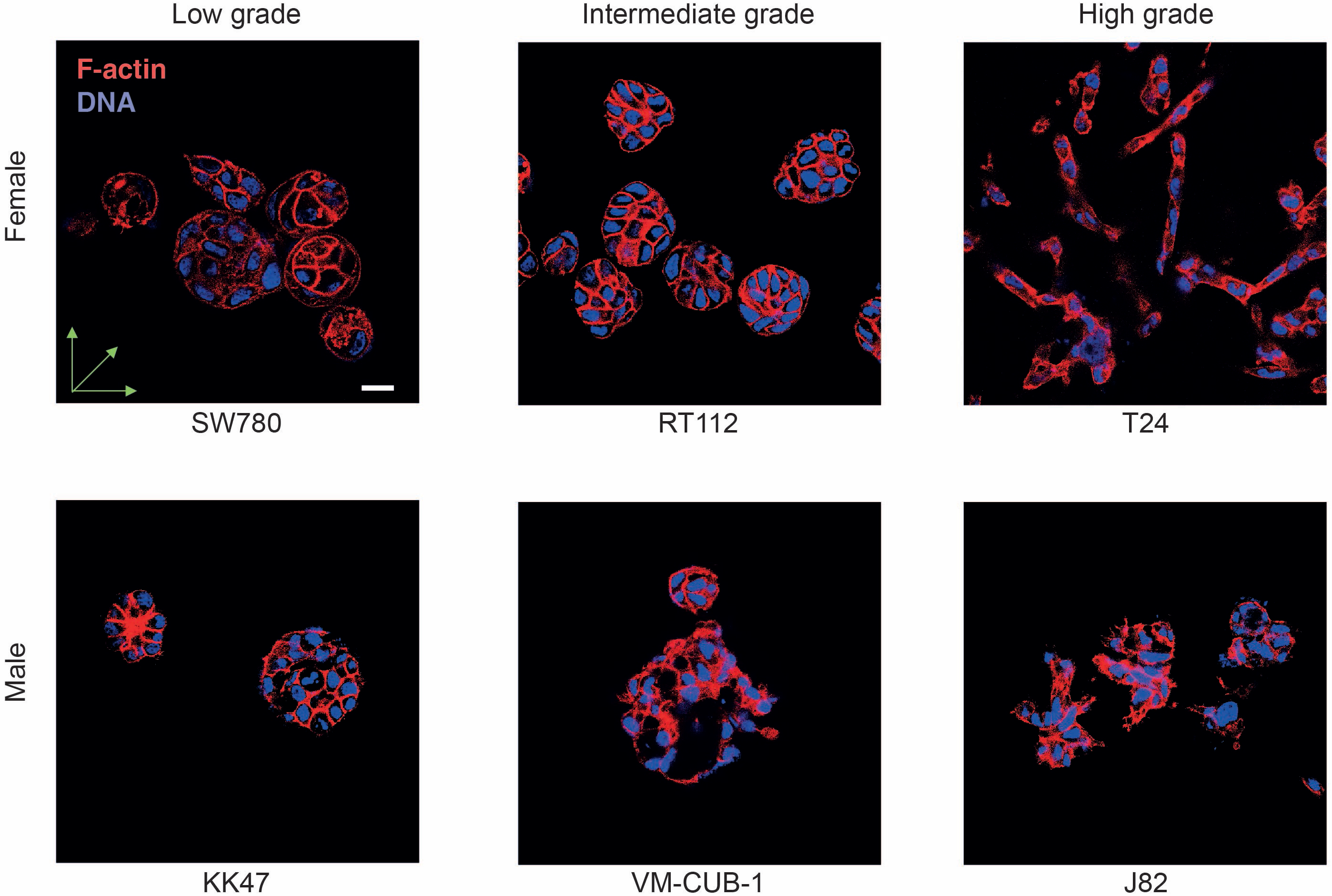
Cell lines derived from patients with high grade bladder carcinoma show greater invasion in laminin-rich scaffolds. Immunofluorescence micrographs of urothelial cancer cell lines SW780 (female, low grade, top right), RT112 (Female, intermediate grade, top middle), T24 (Female, high grade, top left), KK47 (Male, Low grade, bottom left), VM-CUB-1 (Male, intermediate grade, bottom middle) and J82 (Male, high grade, bottom right) cultured within laminin rich scaffolds, stained for DNA (DAPI, blue) and F-actin (Phalloidin, red) and imaged with laser confocal microscopy with maximum intensity projection. Whereas SW780, KK47, RT112 and VM-CUB-1 showed a spherical mass-like architecture, J82 cells exhibited a more invasive grape-like morphology. In comparison, T24 cells showed the highest degree of invasion with stellate multicellular projections in the surrounding matrix. Scale bar, 20 μm).

### Proteomic and bioinformatic analyses reveal elevated levels of Gal-1 in female grade 3 urothelia

To verify our hypothesis, we analyzed and compared the proteome changes across all six above-mentioned bladder carcinoma cell lines. We used gender and grade filters to identify the differentially regulated proteins. The up regulated proteins that associated best with high-grade female-derived cell lines were subjected to a second filter to identify proteins whose genes were known to be responsive to estrogen (33). The top three genes in this list encoded for *ENO1, CLIC4* and *LGALS1* (Fig. 2). The gene encoding GAL-1 not just has an ERE in its near promoter region, it is known to be an estrogen responsive gene (22). Given its cancer-permissive role and its association with epithelial mesenchymal transition (34), we further focused on a potential relation between Galectin-1 and the invasive-proliferative phenotype in female high-grade bladder cells.

**Figure 2:**
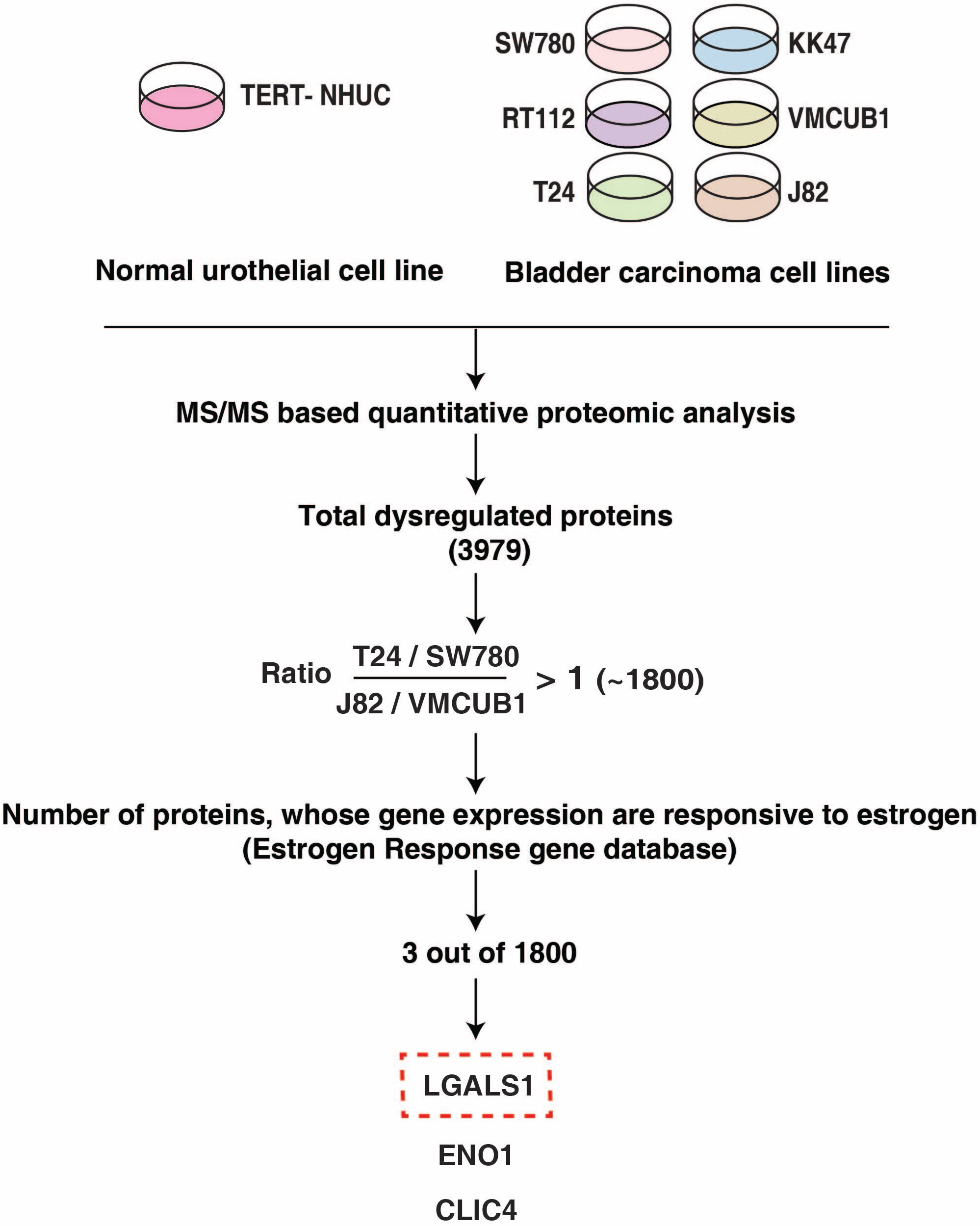
High expression of Gal-1 identified using quantitative proteomics of bladder cancer cell lines. The differential expression of proteins identified in the bladder cancer cell lines compared to non-neoplastic TERT-NHUC cells using LC-MS/MS based approach. Gender and tumor grade-based filters were applied to shortlist the dysregulated proteins and searched in Estrogen Response gene database (ERGDB) to identify ERE in the near promoter region. Gal-1 was observed to be highly expressed in female grade 3 cells.

### GAL-1 mRNA levels associate with bladder cancer cell invasion in 3D

We next sought to investigate a possible association between *LGALS1* gene expression and the morphological phenotypes seen in Fig. 1. We assayed for the levels of *LGALS1* using qPCR and found its mRNA levels to be highest in T24 cells (female grade 3), followed by KK47 (male grade 3), VM-CUB-1 (male grade2), and J82 (male grade 1), and lowest in the epithelial RT112 and SW780 (female grade 1 and 2) (Fig. 3a). We further used indirect immunocytochemistry to observe the localization of GAL-1 protein in the 3D cultures of bladder cancer cells. In high-grade female cancer cells, we observed a dispersed and overall higher signal for GAL-1 in the matrix surrounding the cells (Fig. 3b). This extracellular dispersed staining was absent in lower grade cells (as well as in negative control), although also observed in male high-grade cells. Laminin is a known ligand for GAL-1 (35,36). High gene expression for GAL-1 seems to therefore associate with a distinctly extracellular (in addition to cytoplasmic as noted for all the other cells) localization, which also signifies its well-established secretory matricellular functions. An immunohistochemical comparison of GAL-1 staining showed higher signals in high-grade bladder cancer sections relative to low-grade and ‘adjacent-normal’ tissues (Fig. 3c). We therefore chose to investigate whether the inhibition of GAL-1 levels affect bladder tumor development and growth.

**Figure 3:**
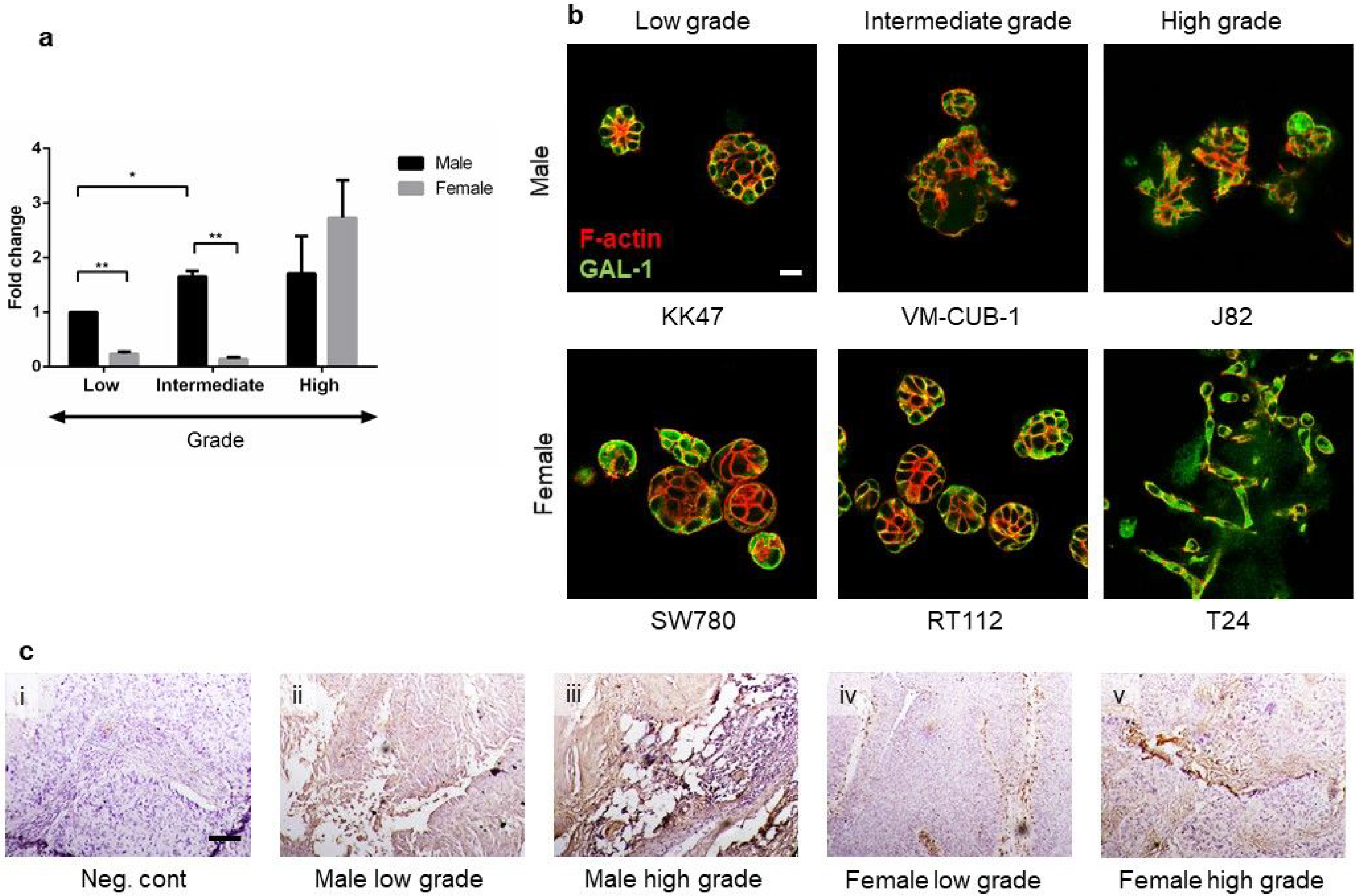
High-grade urothelial carcinoma cells express increased levels of GAL-1. (a)The expression of *LGALS1* gene which encodes GAL-1 when measured across six urothelial cancer cell lines, when assayed using qRT-PCR shows the highest levels in T24 (female high-grade) cells followed by male bladder cancer cells in progressively decreasing grade. In comparison, female intermediate and low-grade cancer cells show lower expression of GAL-1 mRNA. (Internal control: 18S rRNA). (b) Immunofluorescence micrographs of urothelial cancer cell lines KK47 (Male, Low grade, top left), VM-CUB-1 (Male, intermediate grade, top middle) and J82 (Male, high grade, top right) SW780 (female, low grade, bottom right), RT112 (Female, intermediate grade, bottom middle), T24 (Female, high grade, bottom left), cultured within laminin rich scaffolds, stained for DNA (DAPI, blue) and F-actin (Phalloidin, red) and GAL-1 (green) and imaged with laser confocal microscopy with maximum intensity projection, show highest GAL-1 staining in T24 cells with the localization being more extracellular in comparison within with a cellular localization in other cell lines (c) Paraffin embedded tissue sections of bladder cancer stained for GAL-1 (brown; counterstaining with hematoxylin) showed higher expression of GAL-1 in variegated cancer cells within high grade male and female specimen. For all bar graphs, error bars represent SEM. Statistical significance computed using unpaired parametric t-test is given by *P < 0.05; **P < 0.01.

### GAL-1 inhibition decreases the proliferation of high-grade female cancer cells on lrECM

We compared GAL-1 levels across 6 cell lines that were cultured on top of lrECM scaffolds. Using the fluorescence of resorufin formed as a result of the reduction of cell-permeable resazurin as readout, we observed that cells with higher levels of GAL-1 showed greater viability in 3D (Fig. 4a). Pharmacological inhibition of GAL-1 using the calixarene derivative OTX008 (shown to be specific to GAL-1(37,38)) in 3D lrECM scaffolds significantly decreased the ability of T24 cells to form resorufin, suggesting decreased cell proliferation upon attenuation of GAL-1 function (Fig. 4b; dose dependent effect of OTX008 on T24 cell viability shown in Fig. S1). To further confirm this result, we carried out stable knockdown of GAL-1 using lentiviral transduction of T24 cells with shRNA (Fig. 4c; confirmation of knockdown through immunoblotting shown in Fig. S2). We observed that T24 cells with impaired levels of GAL-1 were significantly less proliferative when compared to their control counterparts in 3D.

**Figure 4:**
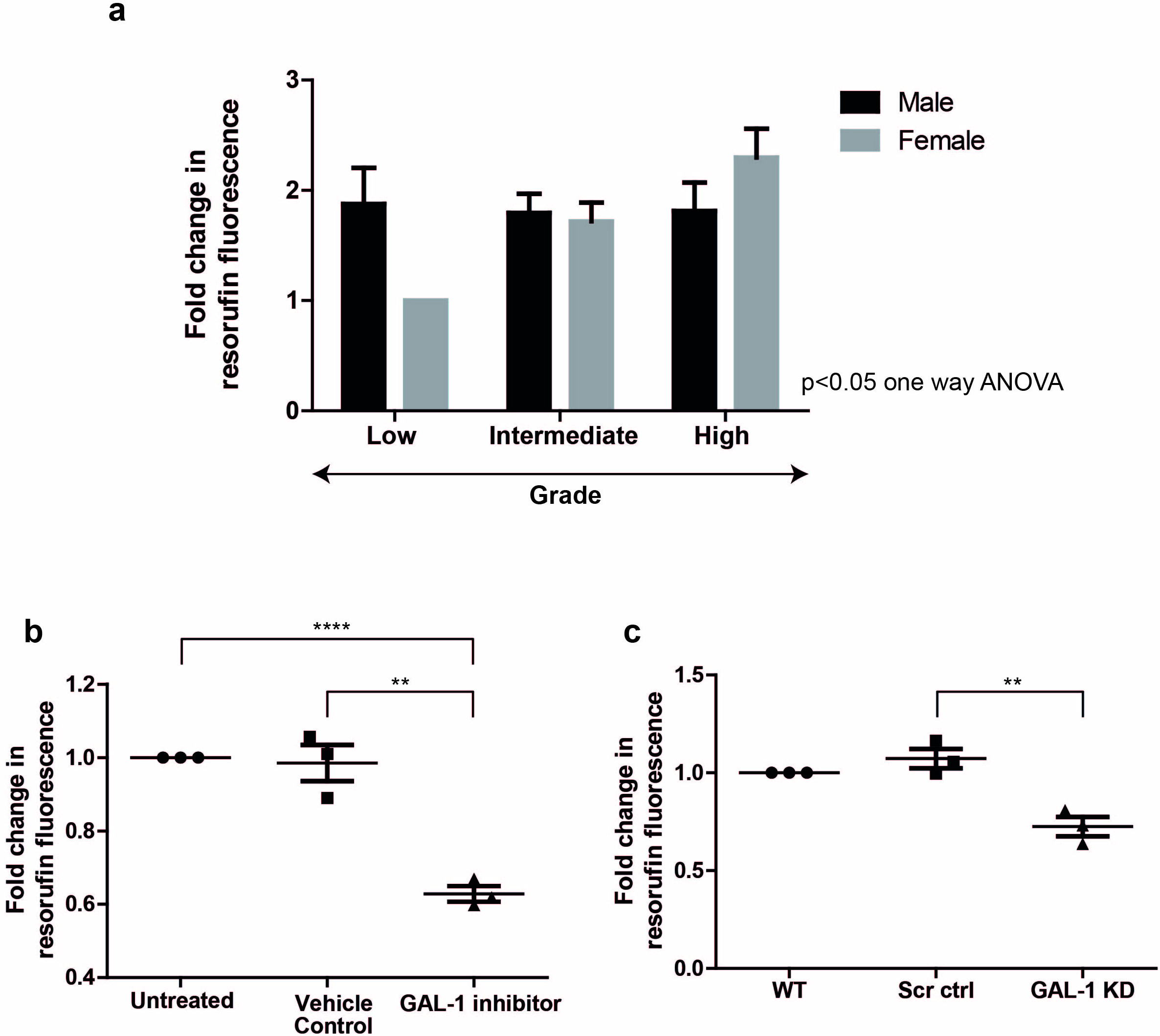
Gal-1 levels promotes proliferation of T24 cells. (a) Among six bladder cancer cell lines cultured on laminin-rich matrix, T24 shows the greatest capacity to convert resazurin to resorufin, a readout of cell viability (n=3). (b) The pharmacological inhibition of GAL-1 by OTX008 in T24 high grade female urothelial cancer cells cultured on laminin-rich matrix (IC_50_: 98.3 μM) leads to decrease in their cell viability (n=3) (c) Knocking down *LGALS1* gene on T24 cells also exhibited decreased cellular viability (n=3), when cultured on laminin-rich matrix (d) For all bar graphs, error bars represent SEM. Statistical significance computed using one-way ANOVA for 4a and unpaired parametric t-test for 4 b and c is given by *P < 0.05; **P < 0.01, ****P < 0.0001.

### GAL-1 inhibition decreases adhesion of invasive bladder cells to laminin

The inability of early grade bladder cancer cells to form multicellular invasions within their matrix surroundings as seen in Fig. 1 can be explained by an increased affinity to form intercellular adhesions and a concomitant decreased affinity for binding to surrounding matrix. To test this hypothesis, we measured the binding of the six bladder carcinoma cell lines to laminin-rich substrata. The adhesion of cells to lrECM was found to be correlated with GAL-1 levels. T24 cells adhered to the greatest extent (Fig. 5a). We tested if adhesion was dependent on GAL-1 expression by adding T24 cells to laminin-rich substrata, with or without treatment with OTX008. We observed that significantly lower number of cells bound to laminin-rich substrata in the presence of OTX008, suggesting that GAL-1 may play a role in helping bladder cells tether better to matrix proteins and facilitate their migration as mesenchymal collectives seen in Fig 1 (Fig. 5b). T24 cells with stable knockdown of GAL-1 showed decreased adhesion to lrECM confirming that GAL-1 could regulate the interaction of cells with laminin-rich milieu (Fig. 5c). An increased ability to tether to matrix commonly accompanies transition of cellular phenotype from an epithelial to a mesenchymal morphology (39). Interestingly, we observed that two cells that showed the greatest expression of GAL-1: T24 and J82, showed no signal for E-cadherin, an epithelial marker whereas N-cadherin was highly upregulated. On the other hand, SW780 and RT112, the cells with low levels of GAL-1 showed high levels of E-cadherin and low levels of N-cadherin (Fig. 5d).

**Figure 5:**
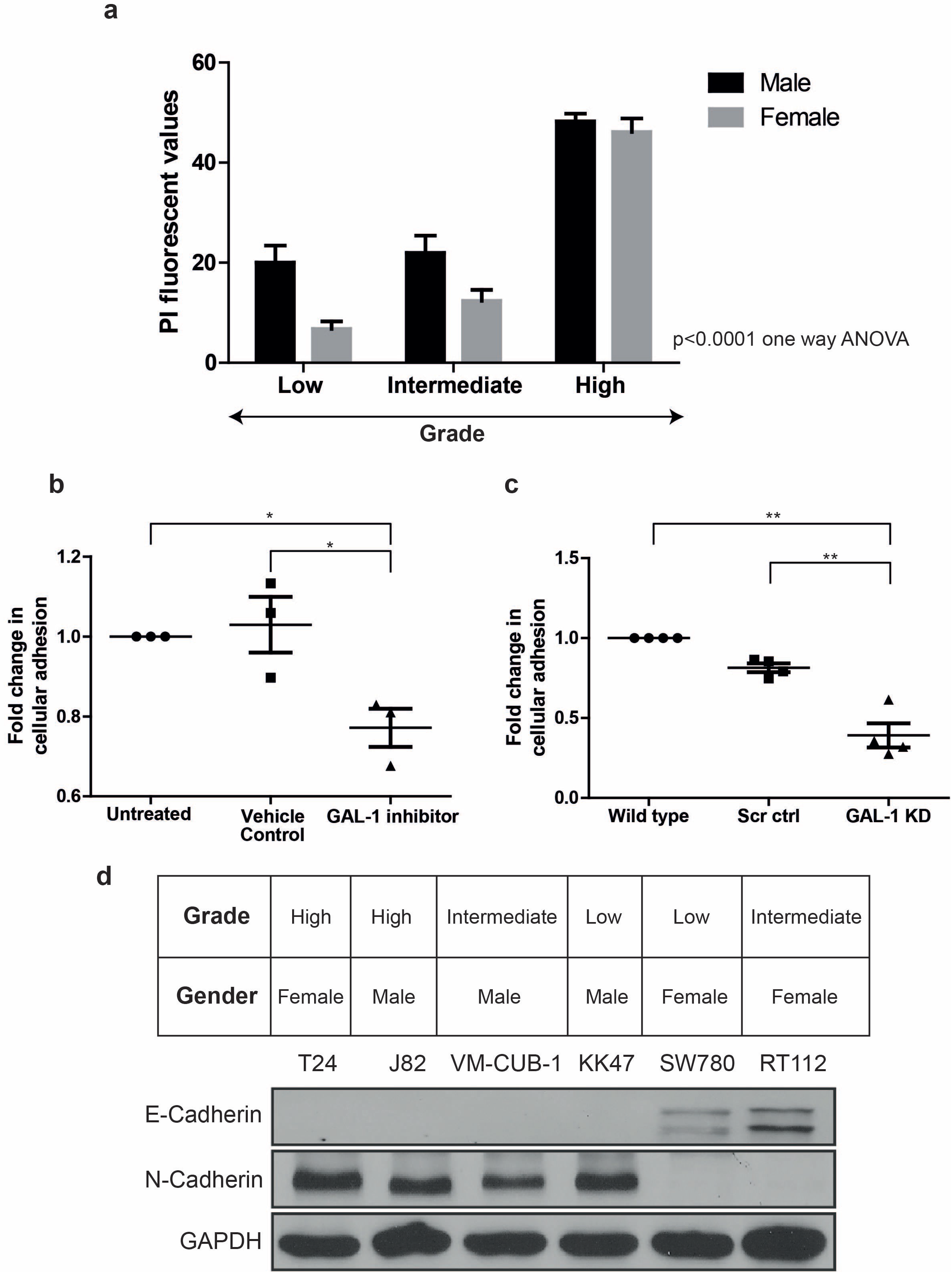
The expression of *LGALS1* gene correlates with the EMT status of urothelial carcinoma cell lines. (a) The adhesion of the six bladder cancer cell lines to lr-ECM measured by their respective PI fluorescence shows highest adhesion by T24 which express highest levels of GAL-1 (n=2). T24 cells (Female, high grade) exhibited decreased adhesion on lr-ECM on (b) sub IC_50_ (50 μM) treatment of OTX008 (n=3) and (c) by knockdown of *LGALS1* gene (n=4). (d) Western blotting of the bladder cancer cells confirmed the expression of E-cadherin (epithelial marker) in SW780 and RT112 (female low and intermediate grade) and highest level of N-cadherin (mesenchymal marker) in T24 (female high grade) followed by male bladder cancer cells, which is an indicative of their respective EMT status. For all bar graphs, error bars represent SEM. Statistical significance computed using one-way ANOVA for 5a and unpaired parametric t-test for 5b and c is given by *P < 0.05; **P < 0.01.

### Inhibition of GAL-1 reverses the invasive phenotype of high-grade female cancer cells

We next sought to investigate the cumulative effect of the regulation by GAL-1 of bladder cancer cell proliferation and matrix adhesion. We probed the effect of GAL-1 expression and function within pathotypic 3D culture assays. The ability of T24 cells to display stellate invasive morphologies when cultured in lrECM typifies the ability of urothelial cancer cells to be able to breach the basement membrane barrier to reach the stromal compartment of the bladder. When we inhibited GAL-1 function within T24 cells, using OTX008, the cells were unable to invasively explore lrECM scaffolds. Rather, they reverted to a spheroid-like multicellular cluster phenotype that is reminiscent of low-grade cellular behavior (Fig. 6a). This was confirmed in T24 cells with stable knockdown of GAL-1, which also exhibited non-invasive morphologies in 3D (Fig 6b).

**Figure 6:**
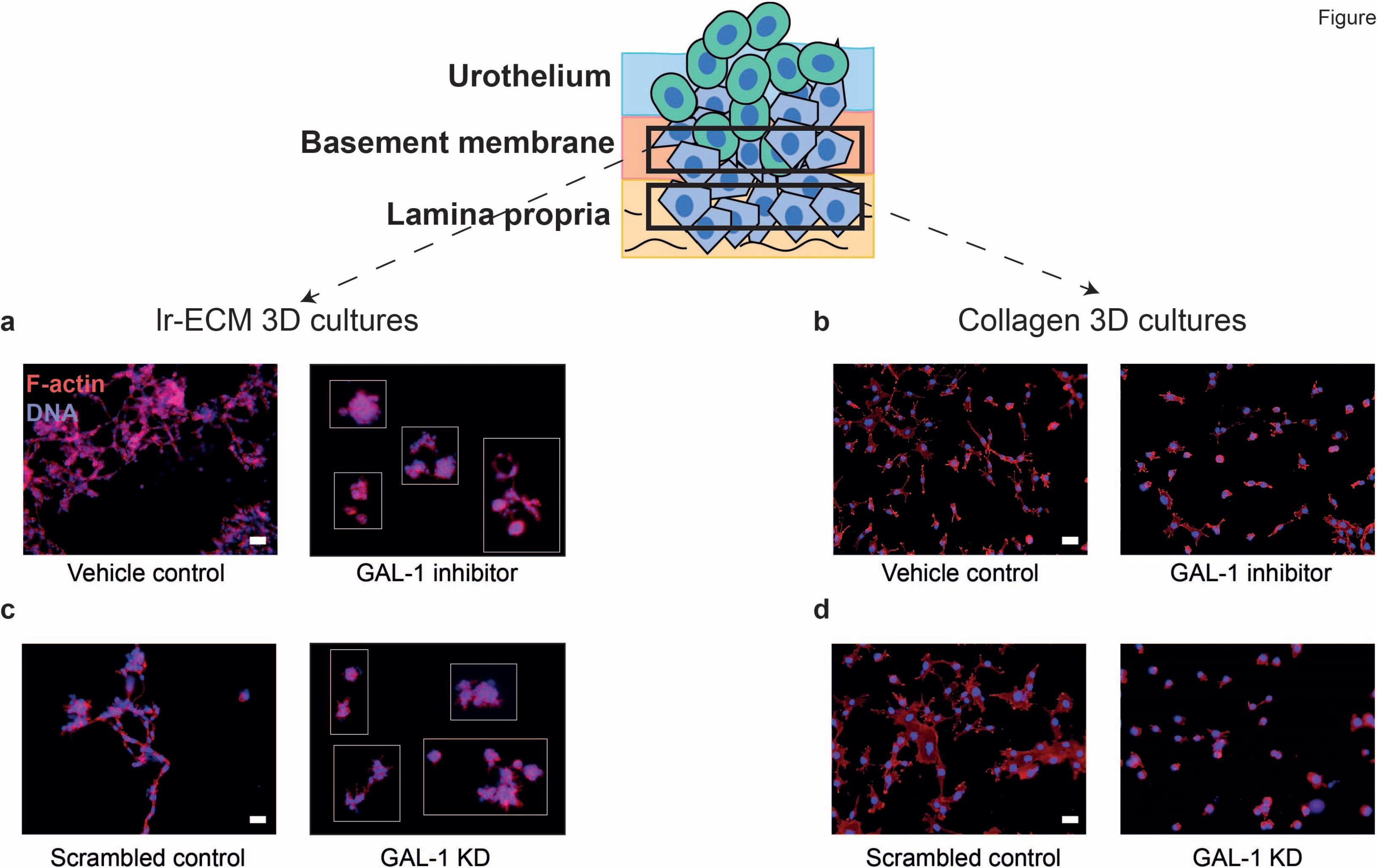
GAL-1 enhances migration of high-grade female bladder cancer cells through laminin- and collagen-rich scaffolds. T24 cells cultured in laminin-rich matrix scaffold migrates through the matrix within 48 hours and forms multicellular invasive trails. Upon treatment with the pharmacological inhibitor of GAL-1 OTX008 at sub IC_50_ concentration (50 μM) T24 cells become rounded and do not invade through their surrounding matrix (a). OTX008 also prevents the invasion of pretreated T24 cells through scaffolds made of the stromal ECM Type 1 collagen (b). T24 cells when transduced with control shRNA exhibit collective invasion within laminin- (c left) and Type 1 collagen-rich matrices (d left). Lentiviral knockdown of GAL-1 in T24 leads to diminished non-invasive phenotype in both scaffolds (c,d right). (Scale bar, 200 μm)

We further tested the role of GAL-1 in migration within stroma, which is rich in Type 1 Collagen. When cells were cultured within Type 1 collagen scaffolds, T24 cells were able to migrate by forming cellular trails through the collagen-rich milieu, ultimately forming a web-like network (Fig. 6c). However, when treated with OTX008, or upon GAL-1 knockdown, the T24 cells remained as single cells embedded within the matrix (Fig. 6c and d).

### GAL-1 regulates the levels of free cell surface sialic acid to mediate bladder cancer cell migration

The regulation of migration and ultimately, metastasis of cancer cells has been linked to cell surface expression of glycans (15,40,41). There is rich literature specifically on sialic acid (Sial), the levels of which have been reported to modulate cell motility impacting invasiveness (12,42,43). When we stained the high-gradeT24 cells for Sialic acid conjugates (Sial) using FITC-conjugated *Limulus polyphemus* agglutinin (LPA), we found that the surface staining of Sial was sparse in comparison with robust levels of in female intermediate-and low-grade cells (Fig. 7a). To our surprise, upon knockdown of GAL-1, Sial levels showed an increase with respect to scrambled and wild type control T24 cells (Fig. 7b and c). We sought to ask whether the effect of GAL-1 levels on Sial levels extended to other cell surface glycans. Upon staining for α1,3-and 1,6-fucose (using *Aleuria Aurantia* lectin), O-linked conjugates (using Jacalin), biantennary N-glycans with bisecting N-acetylglucosamine (GlcNAc) (using Phaseolus vulgaris Erythroagglutinin (PHA-E)), and non-sialylated tri- and tetraantennary N-glycans (using Phaseolus vulgaris Leucoagglutinin (PHA-L)) using flurophore conjugated lectins, we found little difference between glycan levels in control and GAL-1 depleted T24 cells, suggesting our demonstration of the effect of GAL-1 on Sial levels was glycan-specific (Fig. S3). Finally, when control T24 cells were cultured within lrECM and medium was supplemented with sialic acids, the cells showed a decreased tendency for invasive morphogenesis, phenocopying the effect on the invasive phenotype of attenuated GAL-1 function and expression (Fig. 7d)

**Figure 7:**
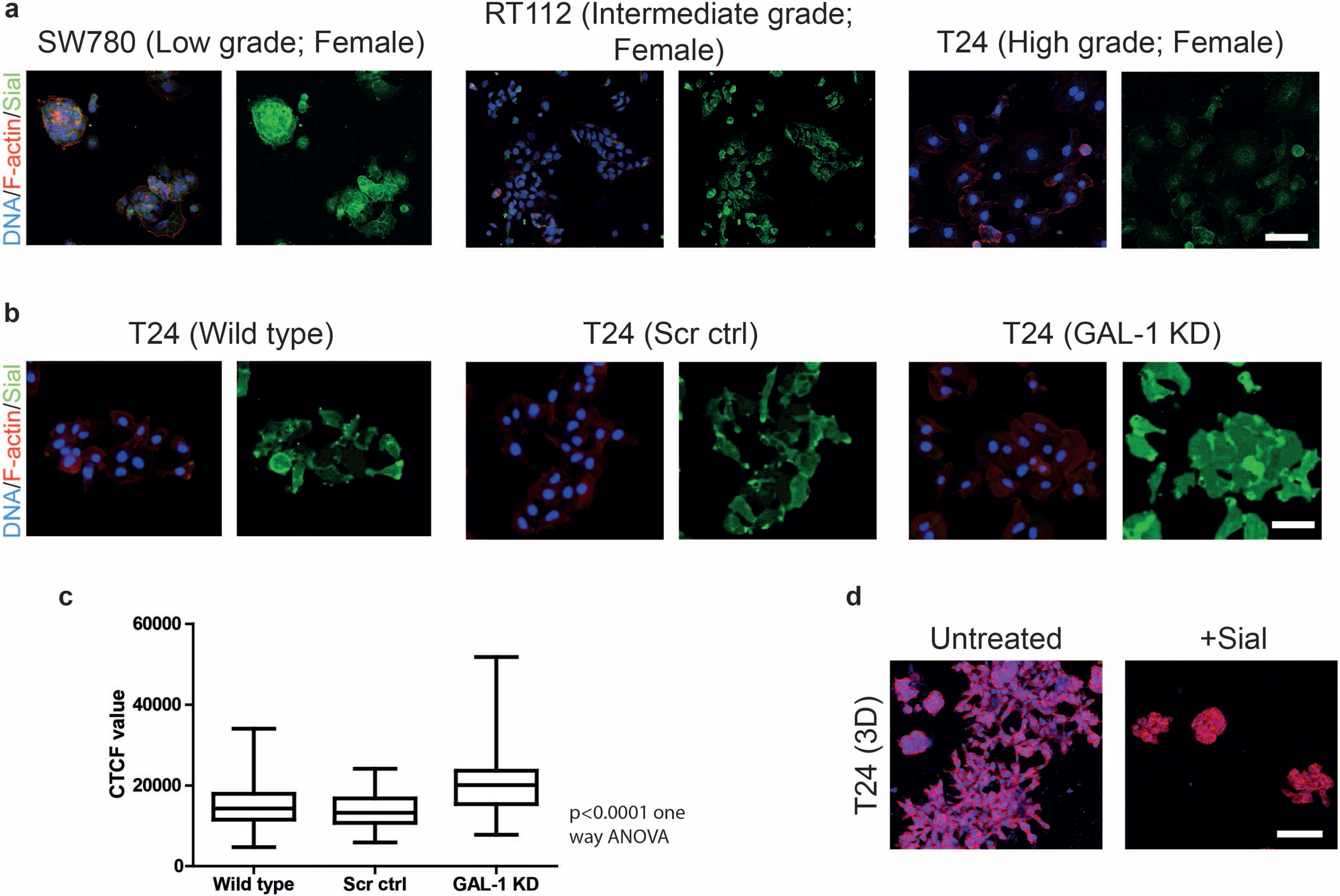
GAL-1 levels correlate with cell surface sialic acid expression in female high-grade bladder cancer cells. (a) Lectin cytochemical staining using *Limulus polyphemus* agglutinin (LPA) showed cell surface staining for sialic acids for female high-grade (T24) bladder cancer cells. In contrast, sialic acid levels were sparse in female intermediate- and low-grade cancer cells. (b) Stable knockdown of *LGALS1* expression in T24 cells resulted in a significant increase in sialic acid levels in T24 cells when compared with levels in scrambled control counterparts. (c) The fluorescent intensity of T24 cells (Wild type, scrambled control and *LGALS1* knockdown) were measured by ImageJ and depicted as box and whiskers plot (d) Upon treatment with exogenous sialic acid (10 mM), the invasive phenotype of T24 cells embedded and cultured within laminin-rich scaffolds (left) reverted to a non-invasive polar phenotype (right) associated with low-grade cancer cells. Statistical significance computed using one-way ANOVA (Scale bar, 100 μm)

## DISCUSSION

The demonstration of the differences between the glycocalyces of normal and cancerous cells are as old as the demonstration that specific genes encode proteins that lead to oncogenesis (oncogenes) or suppression of neoplastic transformation (tumor suppressors) (44,45). Glycocalyces, consisting of glycan structures conjugated to proteins and lipids regulate both the mechano- and chemo-transductive cues within cells (40,41,46). Such glycoconjugates are recognized by- and function in interaction with-, endogenously expressed lectins, of which one of the best-known classes is the β lactoside-binding lectins. It is therefore not surprising that Galectin-1, the first galectin to be characterized, is known to play an important role in cellular homeostasis, neoplastic progression and metastasis in a variety of tissues and organs (47). GAL-1 emerged as a putative candidate in this present study when we applied gender- and grade-based selection filters on a proteomic analysis of a set of six cell lines of low-, intermediate- and high grade, and male and female gender. When we asked which proteins would be highest in the high grade female bladder cancer cell lines, and whose gene expression would be driven by estrogen, GAL-1 was one of the three highest upregulated proteins that satisfied our selection criteria.

In fact, we observed that both the invasiveness of multicellular phenotype in 3D as well as GAL-1 expression shows a nonlinear- and disproportionate- increase with grade in cell lines from females compared with males. The expression of epithelial and mesenchymal markers coincides with the phenotypic spectrum: invasive high-grade female bladder cancer cells showed the high expression of mesenchymal marker, N-Cadherin and a concomitant lack of epithelial E-Cadherin expression. This may provide potentially valuable insights into the observation of how clinically, bladder cancer afflicting women follows a more aggressive trajectory with ill-favored prognosis than in men (3).

A recent study on GAL-1 suggests that it increases the proliferation and invasion of two urothelial cancer lines through upregulation of MMP-9 through Ras-Raf-MAPK-JNK pathway (21). We have built on, and tested, these observations in the context of cell-matrix interactions: proliferation and invasion of bladder cancer cell lines in 3D matrix environments by and large tracks the expression in the cells of GAL-1 mRNA. Notably, the expression of GAL-1 is also congruent with the adhesion of cells to the matrix, which is in keeping with the matricellular roles of GAL-1 (48,49). The female high-grade cells showed higher proliferation, adhesion and invasion within laminin-rich matrix environments. Abrogation of GAL-1 both pharmacologically and using RNAi manages to reverse all these phenotypes. We have examined the subcellular localization of Gal-1: whereas Gal-1 in low grade cancer cells is restricted to the cell surface, it shows a more promiscuous localization in high grade cells with some localization in the nucleus as well as in the extracellular milieu. Such diffuse localization is associated with carcinomatosis (12) and a recent report suggests nuclear Gal-1 binds to and inhibits the tumor suppressive function of Foxp3 (50).

Given our focus on the understanding of the interactions between cancer cells and their microenvironment, we investigated the effect of altering GAL-1 levels on cell surface glycome. Sialic acids which typically are present at the terminal ends of N-and O-linked glycosylation are known to be deregulated in cancers: cancer glycocalyces are typically hypersialylated (42,43). Intriguingly, a report suggests that the 2,6-sialyltransferase ST6GAL1 gets epigenetically inactivated in bladder cancer (51). Our results are in concurrence with the report: we see lower staining of glycocalyx with a lectin that binds to diverse linkages of sialic acids. To our surprise, knocking down GAL-1 results in an increase in sialic acid signals as reported by lectin binding. These results may be interpreted in two ways: GAL-1 may transcriptionally regulate the expression of genes involved in sialic acid biosynthesis. On the other hand, when GAL-1 levels are high and localized extracellularly, they may mask the sialic acids (predominantly 2,3-linked) from binding to LPA resulting in weaker signals from the latter. Our demonstration that addition of free sialic acid (which would not act as a sink for unbound GAL-1) can nevertheless revert the invasive phenotype of T24 cells seems to tilt our opinion towards the first conjecture, i.e., invasion of bladder cancer cells requires lower expression of sialic acids that is mediated endogenously by GAL-1. In the future, we will investigate if GAL-1 may alter Sialic acid levels through promoter methylation of specific sialyltransferases.

Our results suggest that the dysmorphic multicellular phenotype we observed for high-grade bladder cancer could be a cumulative effect of GAL-1 on proliferation, and matrix adhesion of cancer epithelia. The pharmacological and genetic inhibition of Gal-1 was able to arrest the growth and invasion of high-grade invasive cells. It is pertinent to ask if the higher level of GAL-1 is a gender-specific prognosticator of urothelial cancer progression? In order to answer this, we examined TCGA (The cancer Genome Atlas) data stratified on the basis of gender and grades. Whereas high-grade bladder cancer tissues showed significantly higher levels of Gal-1 mRNA in comparison with lower grades, the expression of Gal-1 mRNA in female tissues was not significantly higher than their male counterparts (Fig S4). This could be explained by increase in estrogen levels in male patients with age (52). Therefore, high Gal-1 expression driven by estrogen in carcinomatous urothelia is potentially responsible for the debilitative outcome of the cancer in both men and women but particularly in women. Our functional studies provide a deeper insight into the Gal-1 role in advancing the tumor progression and invasion. Silencing Gal-1 could be a novel therapeutic approach in the management of more advanced bladder cancer in women.

## ACKNOWLEDGMENTS

We would like to thank Jimpi Langthasa for help with preparation of lentiviruses. AB is a recipient of Senior Research Fellowship from Council of Scientific Industrial Research, Government of India. DP would like to acknowledge IISc for his graduate fellowship. KG is a recipient of Senior Research Fellowship from University Grants Commission (UGC), Government of India. PK is a recipient of the Ramanujan Fellowship awarded by Department of Science and Technology (DST), Government of India. RB would like to acknowledge funding support from SERB ECR fellowship (DSTO1586), the DBT-IISc partnership program and the Wellcome Trust Intermediate Alliance Fellowship (WELT0041). In addition, the authors thank the support from the Bio-imaging facility, Division of Biological Science, IISc. The authors also thank Dr. Shoba Krishnappa, Department of Surgical Oncology, Kidwai Cancer Institute, Bangalore, for providing the samples for IHC.

## Supplementary Figures

**Figure S1:** T24 cells treated increasing concentrations of OTX008 to estimate the IC_50_ value. IC_50_ was determined using MTT assay.

**Figure S2:** shRNA-based GAL-1 knockdown in T24 cells. Western blot of GAL-1 knockdown lysates of T24 cells with wild type and scrambled control. α-Tubulin was used as a loading control.

**Figure S3:** Photomicrographs obtained by laser confocal microscopy of T24 cells, both control (top row) and with GAL-1 knockdown (bottom row) stained for α-1,6- and 1,3-linked fucose (using *Aleuria Aurantia* lectin (AAL)), O-linked conjugates (using Jacalin), biantennary N-glycans with bisecting N-acetylglucosamine (GlcNAc) (using *Phaseolus vulgaris* Erythroagglutinin (PHA-E)), and non-sialylated tri- and tetraantennary N-glycans (using *Phaseolus vulgaris* Leucoagglutinin (PHA-L)) (Scale bar, 100 μm)

**Figure S4:** The TPM (Transcript Per Million) value of *LGALS1* gene was obtained from TCGA (The Cancer Genome Atlas) (grade 2 (33 females and 97 males), grade 3 (39 females and 101 males) and grade 4 (35 females and 98 males)) was plotted and the resulting graph depicts significant increase in GAL-1 transcript levels with increase in grades both in female and male patients. Statistical significance computed using one-way ANOVA. For all bar graphs, error bars represent SEM.

## REFERENCES

1. Knowles MA, Hurst CD. Molecular biology of bladder cancer: new insights into pathogenesis and clinical diversity. Nature reviews Cancer 2015;15:25–41

2. Foo J, Leder K, Ryser MD. Multifocality and recurrence risk: a quantitative model of field cancerization. Journal of theoretical biology 2014;355:170–84

3. Dobruch J, Daneshmand S, Fisch M, Lotan Y, Noon AP, Resnick MJ, et al. Gender and Bladder Cancer: A Collaborative Review of Etiology, Biology, and Outcomes. European urology 2016;69:300–10

4. Kumar P, Nandi S, Tan TZ, Ler SG, Chia KS, Lim WY, et al. Highly sensitive and specific novel biomarkers for the diagnosis of transitional bladder carcinoma. Oncotarget 2015;6:13539–49

5. Zhu J, Li Y, Chen C, Ma J, Sun W, Tian Z, et al. NF-kappaB p65 Overexpression Promotes Bladder Cancer Cell Migration via FBW7-Mediated Degradation of RhoGDIalpha Protein. Neoplasia 2017;19:672–83

6. Liang Y, Zhu F, Zhang H, Chen D, Zhang X, Gao Q, et al. Conditional ablation of TGF-beta signaling inhibits tumor progression and invasion in an induced mouse bladder cancer model. Scientific reports 2016;6:29479

7. Chen W, Luo K, Ke Z, Kuai B, He S, Jiang W, et al. TBK1 Promote Bladder Cancer Cell Proliferation and Migration via Akt Signaling. Journal of Cancer 2017;8:1892–9

8. Shang A, Yang M, Shen F, Wang J, Wei J, Wang W, et al. MiR-1-3p Suppresses the Proliferation, Invasion and Migration of Bladder Cancer Cells by Up-Regulating SFRP1 Expression. Cellular physiology and biochemistry: international journal of experimental cellular physiology, biochemistry, and pharmacology 2017;41:1179–88

9. Mao XW, Xiao JQ, Li ZY, Zheng YC, Zhang N. Effects of microRNA-135a on the epithelial-mesenchymal transition, migration and invasion of bladder cancer cells by targeting GSK3beta through the Wnt/beta-catenin signaling pathway. Experimental & molecular medicine 2018;50:e429

10. Friedl P, Alexander S. Cancer invasion and the microenvironment: plasticity and reciprocity. Cell 2011;147:992–1009

11. Stowell SR, Ju T, Cummings RD. Protein glycosylation in cancer. Annual review of pathology 2015;10:473–510

12. Bhat R, Belardi B, Mori H, Kuo P, Tam A, Hines WC, et al. Nuclear repartitioning of galectin-1 by an extracellular glycan switch regulates mammary morphogenesis. Proceedings of the National Academy of Sciences of the United States of America 2016;113:E4820–7

13. Chong Y, Tang D, Xiong Q, Jiang X, Xu C, Huang Y, et al. Galectin-1 from cancer-associated fibroblasts induces epithelial-mesenchymal transition through beta1 integrin-mediated upregulation of Gli1 in gastric cancer. Journal of experimental & clinical cancer research: CR 2016;35:175

14. Chow SN, Chen RJ, Chen CH, Chang TC, Chen LC, Lee WJ, et al. Analysis of protein profiles in human epithelial ovarian cancer tissues by proteomic technology. European journal of gynaecological oncology 2010;31:55–62

15. Laderach DJ, Gentilini LD, Giribaldi L, Delgado VC, Nugnes L, Croci DO, et al. A unique galectin signature in human prostate cancer progression suggests galectin-1 as a key target for treatment of advanced disease. Cancer research 2013;73:86–96

16. Rorive S, Belot N, Decaestecker C, Lefranc F, Gordower L, Micik S, et al. Galectin-1 is highly expressed in human gliomas with relevance for modulation of invasion of tumor astrocytes into the brain parenchyma. Glia 2001;33:241–55

17. Saussez S, Camby I, Toubeau G, Kiss R. Galectins as modulators of tumor progression in head and neck squamous cell carcinomas. Head & neck 2007;29:874–84

18. Paz A, Haklai R, Elad-Sfadia G, Ballan E, Kloog Y. Galectin-1 binds oncogenic H-Ras to mediate Ras membrane anchorage and cell transformation. Oncogene 2001;20:7486–93

19. Vyakarnam A, Dagher SF, Wang JL, Patterson RJ. Evidence for a role for galectin-1 in pre-mRNA splicing. Molecular and cellular biology 1997;17:4730–7

20. Park JW, Voss PG, Grabski S, Wang JL, Patterson RJ. Association of galectin-1 and galectin-3 with Gemin4 in complexes containing the SMN protein. Nucleic acids research 2001;29:3595–602

21. Shen KH, Li CF, Chien LH, Huang CH, Su CC, Liao AC, et al. Role of galectin-1 in urinary bladder urothelial carcinoma cell invasion through the JNK pathway. Cancer science 2016;107:1390–8

22. Than NG, Romero R, Erez O, Weckle A, Tarca AL, Hotra J, et al. Emergence of hormonal and redox regulation of galectin-1 in placental mammals: implication in maternal-fetal immune tolerance. Proceedings of the National Academy of Sciences of the United States of America 2008;105:15819–24

23. Fogh J. Cultivation, characterization, and identification of human tumor cells with emphasis on kidney, testis, and bladder tumors. National Cancer Institute monograph 1978:5–9

24. Marshall CJ, Franks LM, Carbonell AW. Markers of neoplastic transformation in epithelial cell lines derived from human carcinomas. Journal of the National Cancer Institute 1977;58:1743–51

25. Bubenik J, Baresova M, Viklicky V, Jakoubkova J, Sainerova H, Donner J. Established cell line of urinary bladder carcinoma (T24) containing tumour-specific antigen. International journal of cancer 1973;11:765–73

26. Masters JR, Hepburn PJ, Walker L, Highman WJ, Trejdosiewicz LK, Povey S, et al. Tissue culture model of transitional cell carcinoma: characterization of twenty-two human urothelial cell lines. Cancer research 1986;46:3630–6

27. Fanning P, Bulovas K, Saini KS, Libertino JA, Joyce AD, Summerhayes IC. Elevated expression of pp60c-src in low grade human bladder carcinoma. Cancer research 1992;52:1457–62

28. Hisazumi H, Kanokogi M, Nakajima K, Kobayashi T, Tsukahara K, Naito K, et al. [Established cell line of urinary bladder carcinoma (KK-47): growth,heterotransplantation, microscopic structure and chromosome pattern (author’s transl)]. Nihon Hinyokika Gakkai zasshi The japanese journal of urology 1979;70:485–94

29. Kotoh S, Naito S, Yokomizo A, Kohno K, Kuwano M, Kumazawa J. Enhanced expression of gamma-glutamylcysteine synthetase and glutathione S-transferase genes in cisplatin-resistant bladder cancer cells with multidrug resistance phenotype. The Journal of urology 1997;157:1054–8

30. O’Toole C, Price ZH, Ohnuki Y, Unsgaard B. Ultrastructure, karyology and immunology of a cell line originated from a human transitional-cell carcinoma. British journal of cancer 1978;38:64–76

31. O’Toole CM, Povey S, Hepburn P, Franks LM. Identity of some human bladder cancer cell lines. Nature 1983;301:429–30

32. Kenny PA, Lee GY, Myers CA, Neve RM, Semeiks JR, Spellman PT, et al. The morphologies of breast cancer cell lines in three-dimensional assays correlate with their profiles of gene expression. Molecular oncology 2007;1:84–96

33. Tang S, Han H, Bajic VB. ERGDB: Estrogen Responsive Genes Database. Nucleic acids research 2004;32:D533–6

34. Chong Y, Tang D, Gao J, Jiang X, Xu C, Xiong Q, et al. Galectin-1 induces invasion and the epithelial-mesenchymal transition in human gastric cancer cells via non-canonical activation of the hedgehog signaling pathway. Oncotarget 2016;7:83611–26

35. Zhou Q, Cummings RD. L-14 lectin recognition of laminin and its promotion of in vitro cell adhesion. Archives of biochemistry and biophysics 1993;300:6–17

36. Zhou Q, Cummings RD. The S-type lectin from calf heart tissue binds selectively to the carbohydrate chains of laminin. Archives of biochemistry and biophysics 1990;281:27–35

37. Astorgues-Xerri L, Riveiro ME, Tijeras-Raballand A, Serova M, Rabinovich GA, Bieche I, et al. OTX008, a selective small-molecule inhibitor of galectin-1, downregulates cancer cell proliferation, invasion and tumour angiogenesis. Eur J Cancer 2014;50:2463–77

38. Zucchetti M, Bonezzi K, Frapolli R, Sala F, Borsotti P, Zangarini M, et al. Pharmacokinetics and antineoplastic activity of galectin-1-targeting OTX008 in combination with sunitinib. Cancer chemotherapy and pharmacology 2013;72:879–87

39. Kumar S, Das A, Sen S. Extracellular matrix density promotes EMT by weakening cellcell adhesions. Molecular bioSystems 2014;10:838–50

40. Paszek MJ, DuFort CC, Rossier O, Bainer R, Mouw JK, Godula K, et al. The cancer glycocalyx mechanically primes integrin-mediated growth and survival. Nature 2014;511:319–25

41. Zhao Y, Sato Y, Isaji T, Fukuda T, Matsumoto A, Miyoshi E, et al. Branched N-glycans regulate the biological functions of integrins and cadherins. The FEBS journal 2008;275:1939–48

42. Bull C, Stoel MA, den Brok MH, Adema GJ. Sialic acids sweeten a tumor’s life. Cancer research 2014;74:3199–204

43. Seales EC, Shaikh FM, Woodard-Grice AV, Aggarwal P, McBrayer AC, Hennessy KM, et al. A protein kinase C/Ras/ERK signaling pathway activates myeloid fibronectin receptors by altering beta1 integrin sialylation. The Journal of biological chemistry 2005;280:37610–5

44. Aub JC, Sanford BH, Cote MN. Studies on reactivity of tumor and normal cells to a wheat germ agglutinin. Proceedings of the National Academy of Sciences of the United States of America 1965;54:396–9

45. Aub JC, Sanford BH, Wang LH. Reactions of normal and leukemic cell surfaces to a wheat germ agglutinin. Proceedings of the National Academy of Sciences of the United States of America 1965;54:400–2

46. Woods EC, Kai F, Barnes JM, Pedram K, Pickup MW, Hollander MJ, et al. A bulky glycocalyx fosters metastasis formation by promoting G1 cell cycle progression. eLife 2017; 6

47. Johannes L, Jacob R, Leffler H. Galectins at a glance. Journal of cell science 2018;131

48. Thiemann S, Man JH, Chang MH, Lee B, Baum LG. Galectin-1 regulates tissue exit of specific dendritic cell populations. The Journal of biological chemistry 2015;290:22662–77

49. Tsai MS, Chiang MT, Tsai DL, Yang CW, Hou HS, Li YR, et al. Galectin-1 Restricts Vascular Smooth Muscle Cell Motility Via Modulating Adhesion Force and Focal Adhesion Dynamics. Scientific reports 2018;8:11497

50. Gao Y, Li X, Shu Z, Zhang K, Xue X, Li W, et al. Nuclear galectin-1-FOXP3 interaction dampens the tumor-suppressive properties of FOXP3 in breast cancer. Cell death & disease 2018;9:416

51. Antony P, Rose M, Heidenreich A, Knuchel R, Gaisa NT, Dahl E. Epigenetic inactivation of ST6GAL1 in human bladder cancer. BMC cancer 2014;14:901

52. Greenblatt RB, Oettinger M, Bohler CS. Estrogen-androgen levels in aging men and women: therapeutic considerations. Journal of the American Geriatrics Society 1976;24:173–8

